# Using combined RNA/DNA short read sequencing to investigate allele-specific expression from the inactive X chromosome in human cells

**DOI:** 10.64898/2026.05.21.726886

**Authors:** Rachael Thomas, Michael D. Blower

## Abstract

Many genomic regions exhibit allele-specific expression. This effect is most pronounced in imprinted genes, where one copy of a gene is epigenetically silenced, and the inactive X chromosome of female cells, where almost the entire chromosome is silenced. Allele specific gene expression can have significant effects on human health and is implicated in a wide array of diseases. Research into allele specific expression is most often carried out in mouse models where cross breeding of mouse strains can yield progeny with well characterised haplotypes where parent of origin is known for a huge number of SNPs. The same approach cannot be taken with human data and haplotypes must be assembled using expensive and labour intensive long read sequencing and Hi-C based approaches. Although resolved haplotypes are available for a number of cell lines, allowing accurate measurement of allele-specific gene expression, this type of analysis is inaccessible for non-specialist labs. We demonstrate how to use previously published haplotypes to investigate X linked gene silencing and epigenetic changes. Additionally, in this paper we present a method to exploit the profound difference in expression levels between the two human X chromosomes to assign SNPs in expressed RNA to the active or inactive X chromosome using only short read DNA and RNA sequencing. We demonstrate this technique using sequencing libraries generated in house and sequencing data from publicly available databases including for a cell line with a complex karyotype. In each instance we identified genes that were silenced in each cell line opening them up to further research avenues. This X chromosome haplotyping technique can be applied to any clonally derived human cell line with 2 or more X chromosomes allowing researchers to investigate X linked gene silencing in cell lines already present in their lab rather than in the limited number of cell lines for which a haplotype is available.

## INTRODUCTION

Genomic variants can have a profound impact on the expression of genes in many ways; from altering the chromatin landscape and impacting transcription factor binding to post-transcriptional processes such as splicing and translation (reviewed in Cleary and Seoighe, 2021). Allele specific gene expression (ASGE) is widespread and common across the entire genome ^2^ but is particularly pronounced in imprinted genes, many of which are associated with human disease ^3,4^ and the majority of the X chromosome.

In karyotypically normal human female (XX) cells one copy of the X chromosome is randomly inactivated early in development ^5^. This inactivation is initiated by random expression of the long non-coding RNA XIST from one of the X chromosomes, the presumptive inactive X chromosome (pre-Xi), followed by a battery of epigenetic modifications on this chromosome culminating in near chromosome-wide chromatin compaction and gene silencing ^6–8^. This near total silencing of genes on the inactive X chromosome (Xi) leads to X-linked mRNA being predominantly transcribed from the active X chromosome (Xa). It is important to note that some X-linked genes are incompletely silenced and some completely escape from X chromosome inactivation (XCI), many of these genes have been extensively characterised ^9^. Some X linked genes escape XCI only in particular developmental conditions or cell types ^10–13^ and there is significant variability in the extent of XCI escape both between individuals and between cell types within an individual ^9,14–16^. Imbalance in XCI escape and skewed XCI has been implicated in a number of sex-linked human diseases ^17^ including cancer ^18–23^, autoimmune diseases ^24–26^ and neurodevelopmental disorders ^27,28^ due to the high concentration of immune and neural development genes found on the X chromosome.

Genome wide analyses of human disease candidate genes have largely excluded X-linked genes due to the statistical complexity of comparing the single male (XY) to diploid female (XX) X chromosomes ^29^. Whilst most autosomal genes have two copies present in every cell, X-linked genes can be present as a single copy (in male XY cells) or two copies can be present (in female XX cells), this makes it difficult to investigate the effects of variations in these genes at a population level when compared to autosomal genes which are always present with two copies regardless of the sex of the individual. The problem is exacerbated by the random inactivation of one copy of these X linked genes in females, so each individual female is a chimera and the effects of deleterious or protective alleles is dampened by their expression in only half of the individual’s cells. Several previous studies have used sequencing to analyze differential gene expression and epigenomics on the Xi vs Xa. These studies have used mouse strain crosses to maximize SNPs on the X chromosome and non-random inactivation of one of the parental X chromosomes through inducible Xist expression ^8,24,30–40^. These studies exploit the large number of differential mutations between each parental strain’s X chromosome to identify differential gene expression, histone modification and DNA methylation levels. However, the same experimental design is not possible in human cell lines and the number of naturally occurring SNPs in an individual human is considerably lower than in a deliberately cross bred laboratory mouse. To address questions of XaXi regulation of gene expression in humans, studies have compared male (XY) to female (XX) genetic and epigenetic allelic biases ^41^. These studies suffer from the wide variation in expression levels and epigenetic differences between individuals as well as struggling to account for the impact of the Y chromosome in males. In addition, the human reference genome is a composite sequence from many individuals and does not contain the SNPs present in most individuals or cell lines. To further complicate the matter, even when the genomic DNA sequencing of an individual can show which variants are present in that individual’s personal genome further information is required to phase each of these variants to the maternal or the paternal copy of each chromosome. To address this issue many groups have leveraged long read sequencing as well as Hi-C technologies to phase SNPs into a cohesive haplotype capable of providing this parental genotype information ^42–44^. Benchmarks have been established to assess the efficacy of methods of assigning SNPs to parental genomes but these use well characterised parental genomes as a test run for a pipeline rather than an internal check of each haplotyped genome ^45^. There are many options available for allele specific sequencing analysis but most software is not accessible to non-expert bench scientists without extensive bioinformatics support. In this paper we describe a method to leverage knowledge of mammalian X chromosome biology to easily assess the success of a haplotype of a female genome using packages that are simple to implement without previous experience of allele specific sequence analysis.

This paper and its accompanying workflow describe step by step how to use a published phased haplotype to investigate alterations in allele specific expression from the X chromosome. We also present a workflow to *de novo* find SNPs in a cell line from genomic DNA sequencing and use RNA sequencing to identify genes/SNPs subject to XCI and assign them to the Xa or Xi. Our *de novo* SNP assignment of silenced genes shows strong correlation to those phased using complicated and expensive methods and will enable the use of genomic approaches to investigate XCI in a wider variety of cell lines.

## RESULTS

### Using a published haplotype and in-house sequencing libraries to confidently assign XCI status in RPE1 cells

RPE1 cells are derived from human female retinal pigment epithelial cells immortalised with hTERT, they are largely karyotypically normal, aside from a large duplication of a section of chromosome 10 translocated onto one of the X chromosomes, and have 2 X chromosomes, one of which is inactivated and forms a characteristic Barr body, making these cells ideal for the study of X chromosome inactivation (XCI) in human differentiated somatic cells ^42,46^. We leveraged a previously published complete haplotype of RPE1 as well as several RNA-seq libraries generated in our lab to investigate allele specific gene expression of X-linked RNA in these cells ^47^.

We first used Whole Genome Sequencing (WGS) libraries generated from our own population of RPE1 cells to interrogate a published haplotype for RPE1 (RPE1 Variant Call Format (VCF)) ^42^. We used the Personalised Allele-specific Caller (PAC) ^48^ to count reads overlapping each allele of every SNP present in the RPE1 VCF. PAC must be provided with a VCF containing SNPs assigned to the correct parental chromosome which we manually created using the published VCF file and haplotype assignment file ^43^(see methods). To determine if any alleles exhibited unbalanced representation in DNA sequencing data, which might be indicative of a sequencing bias, we calculated the log2 of the ratio of A+1/B+1 reads (that is reads coming from allele version A and reads coming from allele version B, +1 to avoid dividing by zero) (log2abRatio) shown in Figure 1A. This showed wide variation in the log2(abRatio), unexpected for WGS of karyotypically normal cells as each allele should be present at near equal levels to its partner. We used several filtering steps to reduce the presence of these noisy and potentially erroneous SNPs in our downstream analyses. First, we removed all SNPs with zero read coverage which reduced the number of SNPs in our list from 2,317,439 to 2,173,908 meaning 143,531 SNPs could not be replicated in our independent WGS. This could be due to differences in our independent populations of RPE1s or differences in sequencing depth (Supplemental Figure 1A). We filtered for SNPs with >2 reads for A and >2 reads for B and >5 reads for A+B. This further reduced the SNP count to 1,725,511. There were still many SNPs with log2(abRatio) far from the expected log2(abRatio) of around 0 (Supplemental Figure 1B). We further removed all SNPS whose log2(abRatio) was outside of +/- 1 standard deviation from the mean and SNPs with a read count of 30 or more as these SNPs were outliers from the standard SNP coverage of ∼12. Following these filtering steps, we were left with 1,157,875 SNPs for whom we have high confidence and which are represented at nearly equal levels for each allele (Figure 1B) meaning in total we removed 1,180,947 SNPs from the original VCF (a loss of ∼50% of the original SNPs). We used this list to filter RPE1 RNA-seq for SNPs in which we have high confidence.

**Figure 1:**
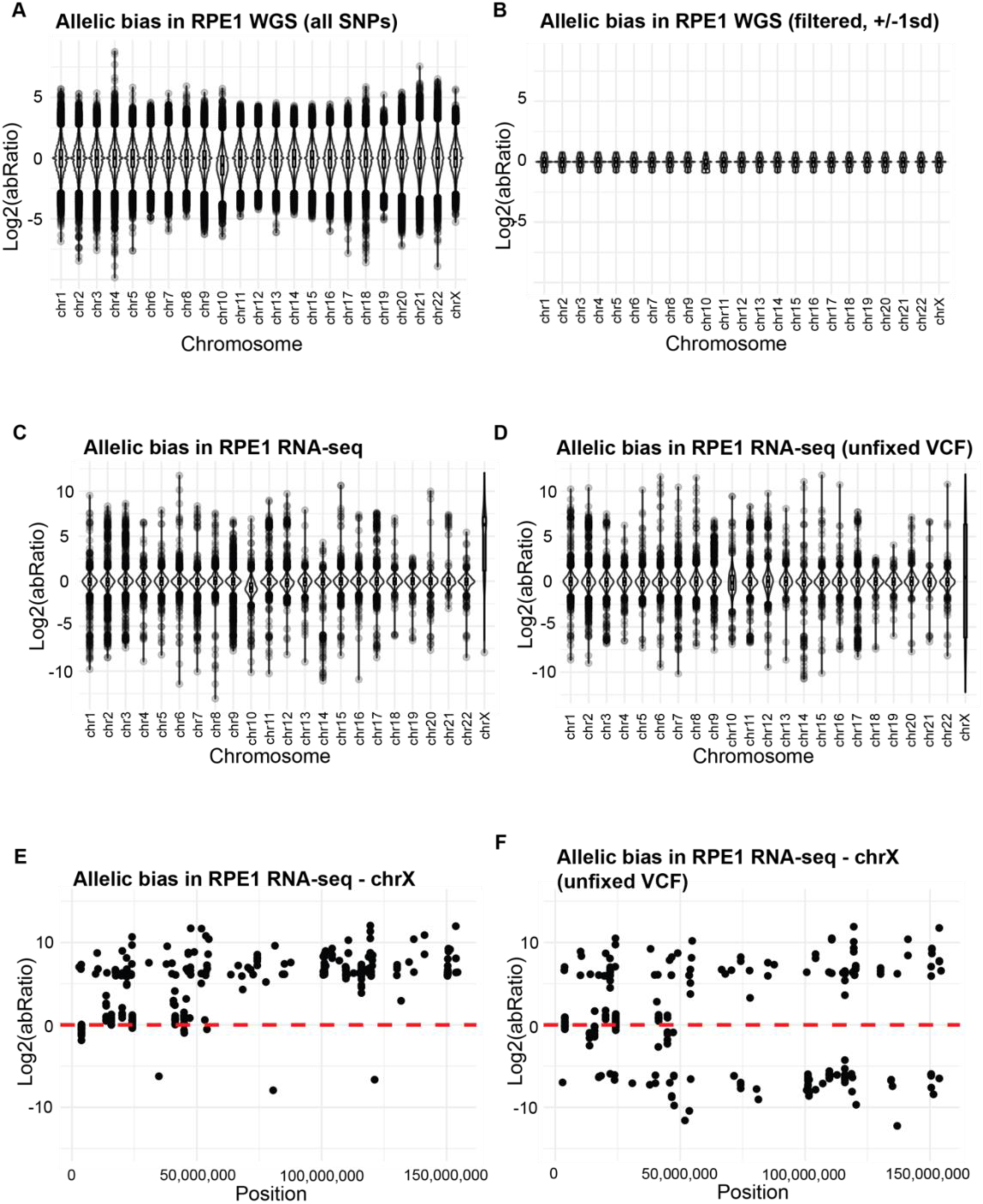
A-D: Plots showing allelic bias (log2(abRatio)) for all chromosome in RPE1 cells from WGS prior to any filtering (A) and post-filtering (B) and from RNA-seq using a filtered and corrected VCF (C) or a VCF where SNPs are randomly assigned (D). E-F: Plots showing allelic bias (log2(abRatio)) in RNA-seq along the X chromosome with a filtered and corrected VCF (E) or a VCF where SNPs are randomly assigned (F).

We additionally noted that there were around half as many SNPs on the X chromosome compared to the autosomes, with an average of one SNP every 2.3-4Kb in autosomes compared to one SNP every 5Kb on the X chromosome. It’s known that the mutation rate is lower on the X chromosome than the autosomes so this is expected ^49,50^.

We leveraged 6 RNA-seq libraries created using our lab population of RPE1 cells to investigate allele specific expression levels using the RPE1 VCF file. Allele specific read counts were summed across libraries and SNPs were filtered for an average of >10 reads per library (>60 reads in total). We then calculated the allelic bias for all SNPs (Supplemental Figure 1C) or SNPs called as high confidence in the WGS (Figure 1C). The allelic bias, when all SNPs are analysed, is highly variable on all chromosomes including a large number of SNPs appearing to be majorly or exclusively expressed from the Xi. When we filtered for high confidence SNPs we found that SNPs located on autosomes had an average log2(abRatio) approaching 0, while the majority of SNPs originating from the X chromosome show a clear bias towards expression from the A allele over the B allele. From this we can assign SNPs of the A X chromosome to the Xa whilst the B allele originate from the Xi. In our dataset we can find 3 SNPs out of 297 informative, high-confidence X chromosome SNPs where reads are assigned to the B allele and none to the A allele, implying they are only expressed from the inactive and not the active X chromosome. Each of these SNPs lie in genomic loci that are not annotated as expressed genes and have high levels of sequence homology with many other areas of the genome and therefore may have been misassigned to the X chromosome however they may represent true Xi biased expression (Supplemental Table 1). This shows the importance of manually verifying the legitimacy of SNP bias before drawing biological conclusions.

We also performed allelic bias analysis for the same RNA-seq libraries but using PAC with the unswapped version of the RPE1 VCF file (i.e random assignment of alleles to parental chromosomes). Figure 1D shows that there is a wide spread of log2(abRatio) values on the X chromosome in the unswapped version, this demonstrates how X chromosome allelic bias would appear in an incorrectly haplotyped VCF. We then analysed allelic bias by position on the X chromosome in correct and randomized VCF files. In contrast to the strong A allele bias in the haplotyped VCF (Figure 1E), analysis of the randomized VCF file shows an almost mirror image of allelic bias along y=0 (Figure 1F).

Interestingly, RPE1 is known to contain a duplication of a large region from chromosome 10 ^42,46^, we observe a number of differential SNPs on chromosome 10 that are more highly expressed from one allele than the other indicating that this region was duplicated onto the Xa (Supplemental Figure 1D).

We have also used this VCF and pipeline to analyse allele-specific differences in histone modifications and chromatin accessibility using Cut&Run and ATAC-Seq data ^47^. We have however found that assays that yield low read counts/coverage often are not able to reliably analyse bias in allele specific effects and this pipeline relies on high coverage to separate signal from noise.

### Analysing allele specific CpG methylation and Xi specific CpG demethylation following 5-Aza-Deoxycytidine treatment

The Xi is known to be hypermethylated at CpG islands (CGIs) compared to the Xa^51^. Bisulfite conversion of genomic DNA followed by sequencing can identify the methylation status of individual CpG islands. Deep sequencing of the whole genome, or even reduced representation bisulfite sequencing (RRBS), would generate a huge amount of data that would be irrelevant to our purposes and therefore costly and wasteful. To address this we used our RPE1 VCF and allele specific analysis of RNA-seq to identify those few X-linked genes that contain differential SNPs in their CGI and are subject to biased expression (see methods). We designed PCR amplicons covering these CGIs and encompassing differential SNPs. Using bisulfite conversion followed by PCR amplification and Oxford Nanopore Sequencing we analysed the methylation status of CpGs within these amplicons, phased to their SNP tag. This allowed for a cost effective, high sequencing depth assay to investigate differences in methylation state of X-linked genes between the Xa and Xi. An example of this methodology demonstrates higher levels of 5mC on the Xi allele of the silenced X linked gene MAMLD1 (Figure 2A). We also observed Xi specific loss of 5mC following treatment of these cells with 5-Aza-2’-Deoxycytidine (Figure 2B).

**Figure 2:**
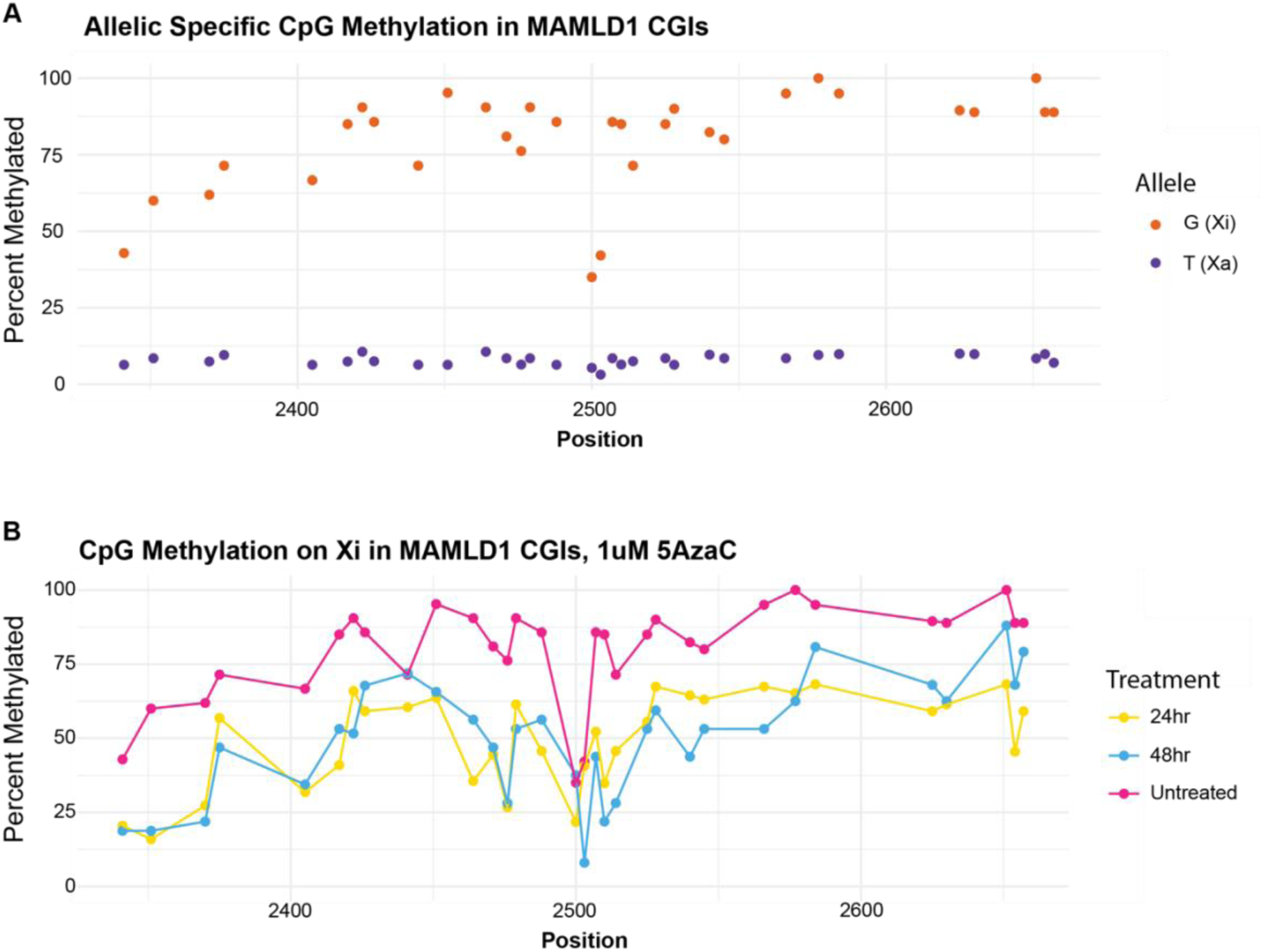
A: Allele specific methylation of CpGs in the CGI of MAMLD1 in RPE1 cells showing higher methylation on the Xi vs Xa. B: Loss of methylation on the Xi allele of MAMLD1 following 24 and 48 hour treatment with 5-Aza-2’-deoxycytodine.

### Identification of derepressed X genes in XIST deleted cells

One practical application of this pipeline is in the identification of desilenced genes following perturbation of XCI factors. One recent study by Bhangu et al introduced somatic deletions of the lncRNA XIST, responsible for the establishment of XCI to investigate derepression of X linked genes following loss of XIST in normal, differentiated cells^52^. To find polymorphic sites they utilised the same haplotype data as in this study, masked those sites whilst aligning reads and then manually counted reads of each genotype that piled up over each polymorphic site of interest. They identified strongly significant changes in expression of three genes following deletion of XIST using this method. We identified the same three genes (MED14, USP9X and DDX3X) as significantly increasing expression from the inactive X chromosome in XIST delete cells vs WT (Supplemental figure 1E). We show that we are able to accurately identify X linked genes that escape from XCI following perturbation of an XCI factor without the need for laborious manual counting using a rapid and largely automated analysis pipeline.

### Using a published haplotype and published sequencing libraries to confidently assign XCI status in IMR90 cells

To determine if we could expand the usage of this pipeline to additional cell lines and publicly accessible datasets we used a VCF file (IMR90 VCF) for the human female immortalised cell line IMR90 ^53^. We followed the same initial preprocessing steps to manually swap alleles within IMR90 VCF. We counted allele specific reads in WGS data from [SRX7735728] using PAC to identify a list of high-confidence SNPs, removing a total of 1,662,854 SNPs for a final SNP count of 546,628 SNPs (Figure 3A and 3B). We used this swapped and filtered VCF file (IMR90 filtered VCF) to run PAC on RNA-seq data from [SRX24925913 and SRX24925914] to assess allele specific expression of genes and assign the Xa and Xi. We filtered for >10 reads covering the SNP as we found that low coverage SNPs could not be confidently assign allelic bias. We observed a clear bias towards reads originating from one chromosome and could confidently assign this chromosome as the Xa. The need for removal of the low confidence, low coverage SNPs is illustrated in Figure 3 C-F and shows the importance of deep sequencing to confidently assign bias to a SNP. Figure 3 C and D show the allele specific bias of SNPs that passed the filtering steps, we can see a clear bias towards the “A” allele in expression that is uniform along the length of the X chromosome. Prior to filtering (Figure 3 E and F) we observe many SNPs with a strong bias towards the “B” allele, the majority of these had a very low read coverage. This work shows that our pipeline can be applied to publicly available datasets and was not specific to our in-house sequencing libraries.

**Figure 3:**
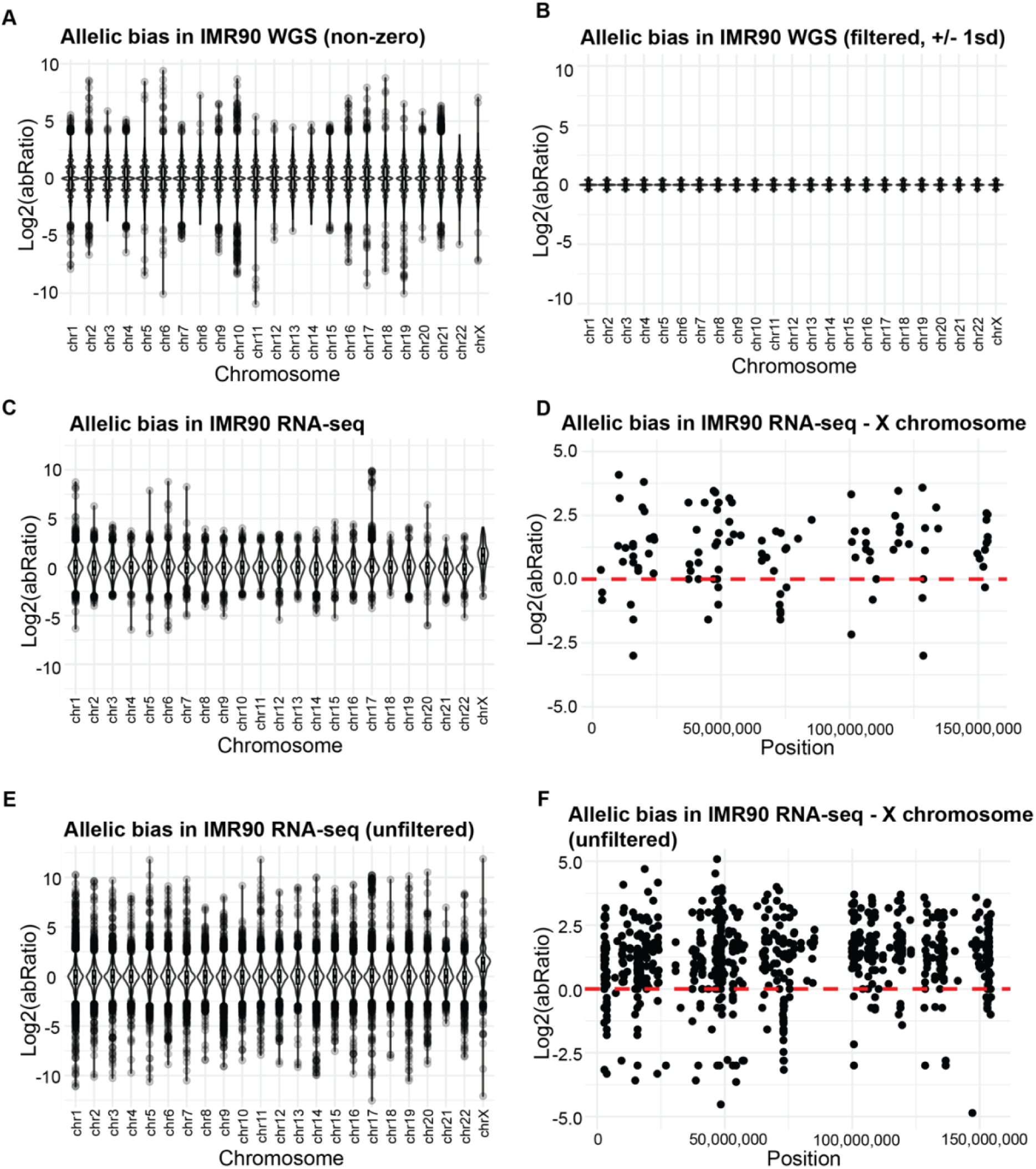
A-B: Plots showing allele specific bias in WGS of IMR90 cells pre-filtering (A) and post-filtering (B). C-F: Allele specific bias in RNA-seq of IMR90 cells for all chromosomes (C and E) or the X chromosome only (D and F) either using a filtered set of SNPs (C and D) or all SNPs (E and F).

### Assigning SNPs in XCI silenced genes to the active or inactive X chromosome using only short read sequencing

Whilst some female cell lines of interest have published haplotypes available, many do not. We investigated whether we could use short read illumina sequencing combined with our understanding of X chromosome biology to replicate the assignment of SNPs to the Xa or Xi that was produced using a laborious haplotype assignment. We used GATK to generate de novo unphased haplotypes for the X chromosome using our WGS library from our population of RPE1 cells^54^. We used this unphased VCF to count reads in our six RNA-seq libraries to identify SNPs with high levels of biased expression (indicative of genes subject to XCI) (Figure 4A). We observed a population of SNPs with log2(abRatio) close to 0, indicative of SNPs not subject to XCI silencing. We could also see two “lobes” of SNPs indicating pronounced bias towards either the “A” or “B” allele, as expected for unphased, and therefore random, haplotypes. Using this information, we manually switched the chromosome of origin for all XCI subject SNPs and artificially “phased” them to the same chromosome with all SNPs showing log2(abRatio) bias in the same direction (Figure 4B). SNPs that could not be confidently assigned, as they had a log2(abRatio) close to 0 were excluded from further analysis. We compared silenced SNPs discovered in this way (446 SNPs) to silenced SNPs from the RPE1 VCF^42^ (229 SNPs) and found 222 SNPs in common. The log2(abRatio) of these 222 SNPs in common was found to be highly correlated between the two VCF files used (Spearman’s Rank Correlation 0.997, Figure 4c). This demonstrates that when searching for genes subject to XCI in a cell line it is not essential to perform costly and complicated sequencing experiments and simple, short read DNA and RNA sequencing is sufficient to generate a list of genes of interest for further study.

**Figure 4:**
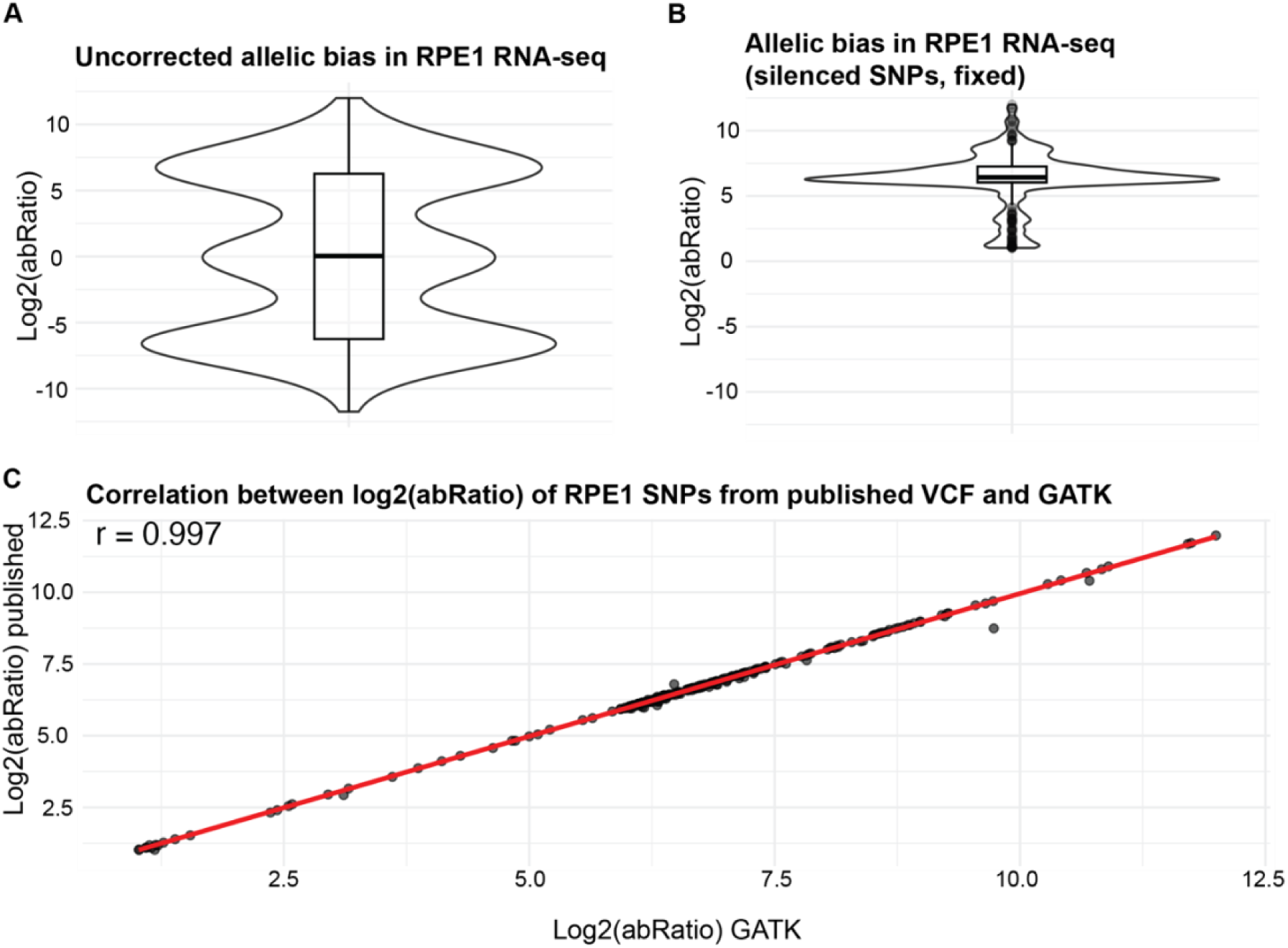
A-B: Allelic bias from RNA-seq of RPE1s in SNPs identified using GATK, all SNPs (A) or SNPs displaying clear bias (-1 > log2(abRatio) > 1) and with all negative bias transposed to positive (B). C: Correlation between allelic bias in SNPs identified using published haplotype data and SNPs identified using GATK.

### Assigning SNPs in XCI silenced genes to the active or inactive X chromosome using only short read sequencing in complex karyotype

HEK293T cells are another popular immortal cell line of female origin. HEK293T cells are a highly transfectable derivative of HEK293 human embryonic kidney cells and, unlike the karyotypically normal RPE1 and IMR90 cell lines, HEK293T cells are described as hypertriploid and display varying ploidy throughout any population. Of particular note HEK293Ts have been observed to contain 2 distinct Barr bodies indicating at least 2 inactive X chromosomes. Karyograms of HEK293 cells (the parental cell line of HEK293T) indicate at least 3 full X chromosomes as well as additional X chromosome material and confirm that total chromosome numbers are variable within a population ^55^. To determine if we could identify Xa SNPs de novo within this complex karyotype using the approach described above we used GATK to generate de novo unphased haplotypes from WGS of HEK293T assuming approximately triploid coverage of the X chromosome ^56^. As we would expect, unlike with the diploid RPE1, the log2(abRatio) of these SNPs did not trend towards 0. We would expect strictly triploid cells to trend towards 1 or -1 however even in our high coverage SNP loci the log2(abRatio) ranged within approximately +/- 3 indicative of a copy number above 3 in many instances (Figure 5A). This reflects the unstable ploidy and additional X chromosome material typical of HEK293T cells. We counted allele specific reads with PAC and used log2(abRatio) to manually swap alleles as described above (Figure 5B and 5C). This allowed us to generate a list of 381 SNPs subject to XCI silencing and identify whether they are present on the active X chromosome. We also investigated whether the allelic bias in RNA expression could be explained by copy number variation at that SNP but found that there was no correlation between log2(abRatio) from WGS of a SNP and log2(abRatio) from RNA-seq (Figure 5D) indicating that additional copies of that allele in the genome did not account for increased expression of that allele in the transcriptome.

**Figure 5:**
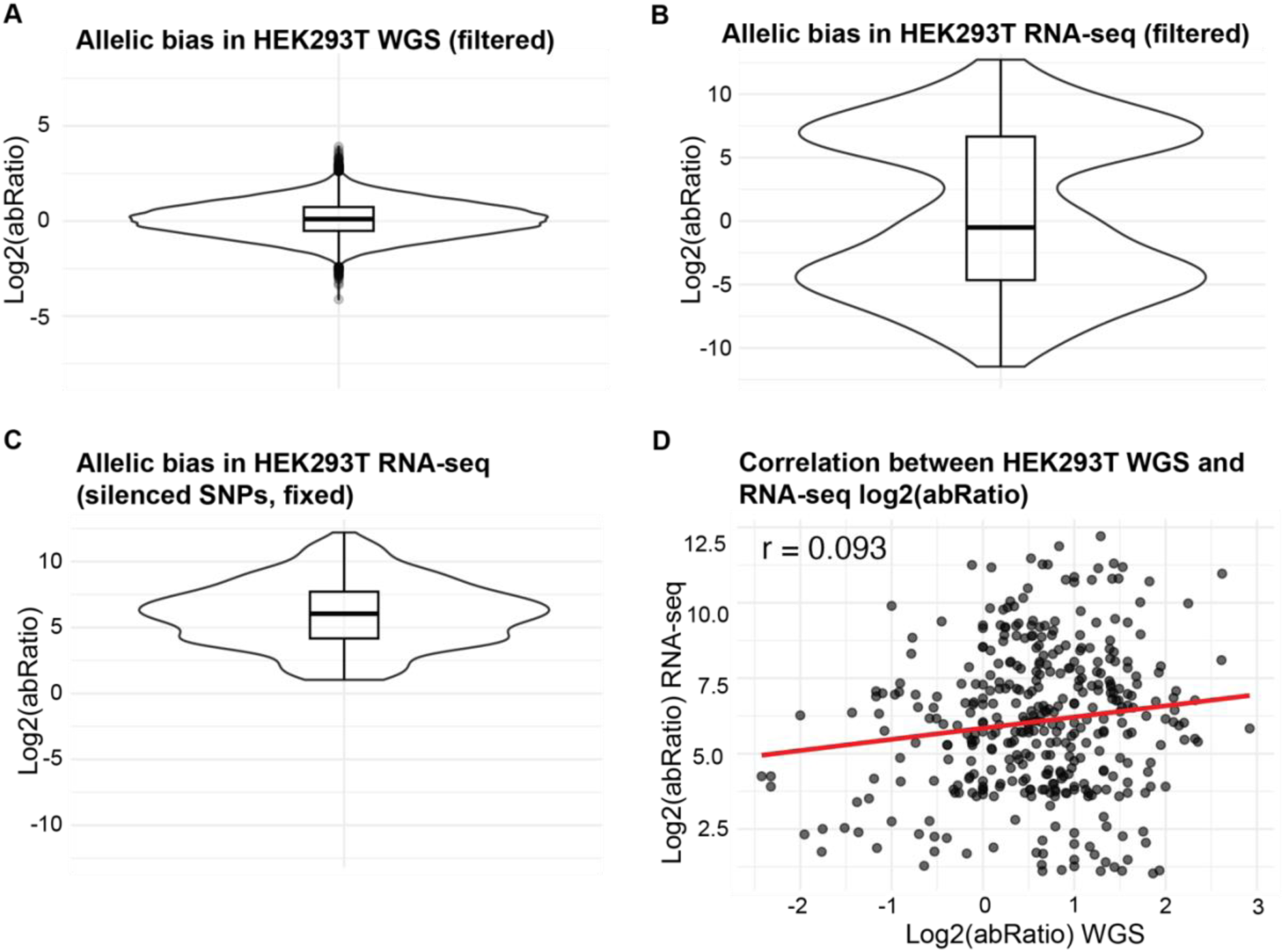
A: Allelic bias in WGS of HEK293T. B-C: Allelic bias from RNA-seq of HEK293T in SNPs identified using GATK, all SNPs (B) or SNPs displaying clear bias (-1 > log2(abRatio) > 1) and with all negative bias transposed to positive (C). D: Correlation between allelic bias of SNPs in WGS and in RNA-seq.

### XCI status cannot be confidently assigned to mixed population cells

It is important to note that the same pipeline described above cannot be applied to available sequencing from all human female sources. For instance, there is a published haplotype-resolved VCF file from a human female individual from whom many tissues and generated cell lines have been sequenced ^57^. After following the above pipelines, when analysing RNA-seq of cultured lung cells from this individual, even the most stringent filtering could not confidently assign either X chromosome as active or inactive (Supplemental Figure 1F). Whilst this could be due to errors within the VCF file this may instead indicate that the cultured cells were not derived from a clonal population. As XCI happens early in development and is faithfully maintained through all subsequent cell divisions, cells taken from one individual will have a mixture of cells with the X chromosome from each parent of origin inactivated. As information is not available about whether this cell line was generated from a single cell isolate we cannot make any conclusions. It is therefore important to consider the experimental design of publicly available datasets before attempting to apply this pipeline.

### X-linked genes silenced in common between RPE1, IMR90 and HEK293T

There is large variation in which genes remain silenced between individuals and even between cell types within the same individual^15,16^. We investigated genes that were silenced in all 3 cell lines where we could confidently assign expression (Figure 6A). We found that of the total 239 silenced genes containing SNPs in any of the 3 cell lines, only 3 silenced genes were in common across all 3 cell lines (MID1, MBTPS2, ACOT9), this is just over 1% of the total gene list. Of the 93 genes silenced in RPE1, 46 of them are in common with HEK293T (which has 178 silenced genes). To investigate whether this low number of genes in common is due to very few genes containing SNPs in common between the cell lines we carried out the same analysis using SNPs found in WGS rather than found to be silenced in RNA-seq. We found more genes containing SNPs in our WGS datasets compared to the RNA-seq datasets for all cell lines, this may be because these genes are not expressed in these cell lines to a level where they would be excluded from our RNA-seq analysis. We found of the 779 genes containing SNPs across the 3 cell lines, 366 genes were in common between all cell lines (47%) (Figure 6B). The low number of silenced genes in common between the 3 cell lines may be due to differences in cellular specialisation (RPE1s originate from retinal epithelium, HEK293Ts from embryonic kidney and IMR90s from embryonic lung) and therefore cell specific gene expression

**Figure 6:**
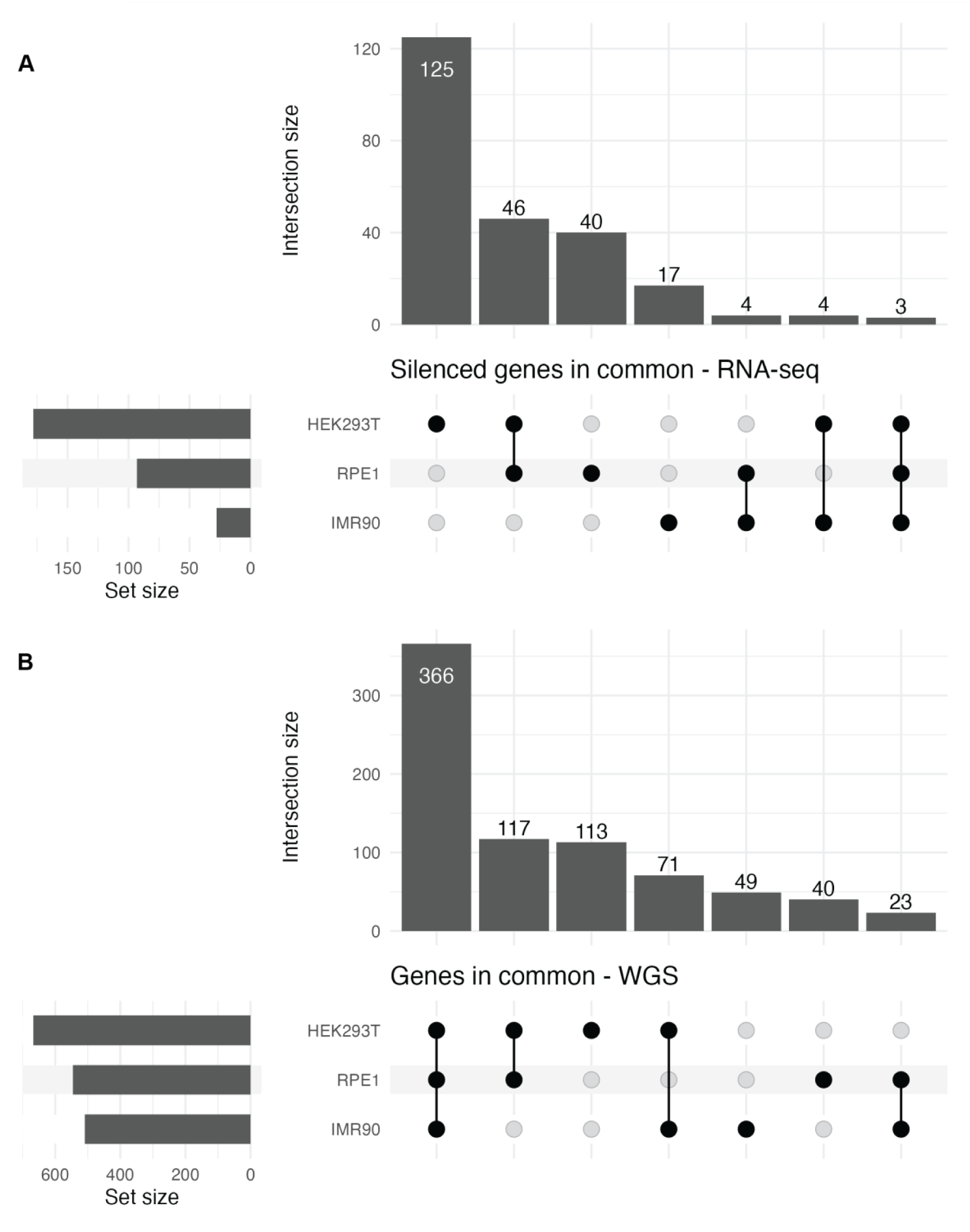
Upset plots showing the intersection of genes containing SNPs and identified as silenced in RNA-seq from RPE1, HEK293T and IMR90 (A) or all genes containing SNPs identified in WGS (B).

## METHODS

### Whole Genome Sequencing

Genomic DNA was extracted from RPE1 whole cells using the One-4-All Genomic DNA Miniprep Kit from Bio Basic (BS88505). Sequencing libraries were prepared using the NEBNext Ultra II DNA Library Prep Kit for Illumina from New England Biolabs (E7645S) and sequenced to a depth of 20x at Novogene using paired-end 150bp reads.

### RNA-sequencing

rRNA depleted libraries were generated using NEB Next Ultra II RNA library kit. Libraries were sequenced at Novogene using 150bp paired-end reads.

### Bisulfite-sequencing

RPE1 cells were treated with 1uM 5-Aza-2’-deoxycytodine (Sigma, A3656) (or DMSO) for 24 or 48 hours.

Genomic DNA was extracted from RPE1 cells using the One-4-All Genomic DNA Miniprep Kit from Bio Basic (BS88505). Genomic DNA was bisulfite converted using Zymo Research EZ DNA Methylation-Lightning Kit (D5030). Bisulfite converted DNA was amplified with CG free primers and ZymoTaq Premix (E2003). CG free PCR primers were designed using MethPrimer bisulfite sequencing primer designer ^58^ to encompass as much of the CGI as possible and capture SNPs within their amplicon, and were ordered from Eton Bioscience. Amplicons were sequenced using Oxford Nanopore Technologies amplification-free long-read sequencing by Plasmidsaurus.

### Sequencing analysis steps in common

Fastq files of sequenced libraries (WGS and RNA-seq) were analysed for quality, adaptor sequences were trimmed, and duplicate reads were removed using the fastp software suite ^59^.

Trimmed and de-duplicated reads were passed to the PAC pipeline ^48^ along with the VCF of haplotyped SNPs ^42^ using the Singularity profile and the hg38 genome build. Of note, PAC expects the alleles in the VCF file to already be phased i.e. it will not look for any further columns to indicate whether the randomly assigned A or B was correct. Prior to running PAC we used the hap_monosomy column from the accompanying final haplotype file and used a custom awk script to manually switch the assignment of each SNP in the REF and ALT position. If the genotype was indicated as 1 the SNPs remained in place, if the genotype indicated as -1 then the contents of REF and ALT were swapped. If the haplotype could not be assigned (indicated as 0) the SNP was removed from the VCF.

### PAC calls from in-house WGS to filter SNPs from published haplotypes

Using R we inspected the output read counts overlapping SNPs in our WGS library. We first calculated the log2 of the ratio of A+1/B+1 reads (that is reads coming from allele version A and reads coming from allele version B, +1 to avoid dividing by zero) (log2(abRatio)). We filtered SNPs for >0 total reads, >2x coverage at each allele and >5x coverage at either allele of the SNP and log2(abRatio) was outside of +/- 1 standard deviation from the mean and <40 total reads.

### Analysing PAC called from in house RNA-seq of RPE1

Using R, PAC read counts were summed across 6 libraries and SNPs were filtered for an average of least 10 reads per library (>= 60 reads in total). SNPs that did not appear high confidence in the WGS step were discarded. Log2(abRatio) was calculated as described previously.

### Allele specific methylation analysis of RPE1

X chromosome genes with CpG islands (CGI) that contain differential SNPs were identified by intersecting a bed file of X chromosome CGI regions from UCSC Table Browser (cpgIslandExt, hg38, chrX) with the filtered VCF file (described above) using bedtools ^60^. We manually inspected the location of these CGI SNPs and assigned them to their closest genes and filtered for genes highly expressed in our RPE1 population and subject to XCI silencing.

We processed raw fastq files using Trim Galore, a wrapper for Cutadapt and FastQC for consistent adaptor and quality trimming suited for reduced representation bisulfite sequencing (RRBS) data. Custom parental genomes were generated for each CGI using samtools faidx to create a FASTA file of just the CGI region. We then used a custom AWK script to find the SNP within the CGI region FASTA and make versions of the FASTA file with each allele at that locus. We then used Bismark’s bismark_genome_preparation to in silico bisulfite convert each allele of the FASTA file ^61^. Fastq files were aligned to each of the bisulfite converted parent genomes using Bismark. We used a custom python script using pysam to split outputted bam files by the allele at each differential SNP and write them to new, allele-specific, bam files. We then used samtools merge to combine allele-specific bams containing the same allele but originating from alignment to different parental genomes. We used samtools collate, fixmate, sort and markdup to remove reads that aligned to both versions of the genome and were present in both bam files. We used bismark_methylation_extractor to extract cytosine methylation information for each read from each allele’s bam file. Methylation levels at each CpG were examined using R and compared between alleles.

### Finding Xa SNPs in publicly accessible data – IMR90

We manually swapped REF and ALT of each SNP according to genotype information for the VCF file for IMR90 ^53^ as described above and ran PAC on WGS data from [SRX7735728]. As described above we used allelic bias (log2(abRatio)) to filter the VCF for high confidence SNPs. We used this swapped and filtered VCF file to run PAC on RNA-seq data from [SRX24925913 and SRX24925914] to assess allele specific expression of genes as described above. Using R we filtered for >10 reads covering the SNP.

### Identifying SNPs de novo from in house WGS – RPE1

Raw fastqs were processed using fastp as described above. Trimmed and deduplicated fastqs were aligned to hg38 using BWA-MEM ^62^. We used samtools to subset the output BAM file for chrX and used GATK to assign read groups (AddorReplaceReadGroups), mark duplicates (MarkDuplicates) and call haplotypes (HaplotypeCaller).

### Identifying SNPs from publicly accessible WGS – HEK293T

Fastq files of WGS of HEK293T were downloaded from [SRX24505563] using sratoolkit and cleaned for paired end reads using SeqKit ^63^. Raw fastqs were processed and haplotypes called as described above. GATK HaplotypeCaller was used with the command –sample-ploidy 3 and -L chrX to account for the approximate triploidy of chrX. PAC requires a diploid haplotype so we artificially “diploidised” the outputted VCF from GATK using bcftools to skip complex genotype (1/2/3) and code “diploid-like” genotypes in an acceptable way for PAC ( i.e. 0/0/1 to 0/1, 0/1/1 to 0/1 and 1/1/1 to 1/1). PAC is very memory intensive so we converted the chrX subsetted BAM file to make chrX only fastq files using samtools. We ran PAC with these chrX only fastq files and our custom VCF. Log2(abRatio) of each differential SNP was analysed using R. We subsetted the SNPs for a coverage of >5 reads per allele and >10 reads in total and used this as our set of high confidence SNPs.

### HEK293T RNA-seq to assign haplotypes

Fastq files for 5 RNA-seq libraries of HEK293T were downloaded from [SRX27138104, SRX27138103, SRX27138102, SRX27229863, SRX27229862] and processed as described for WGS. Trimmed and deduplicated fastqs were aligned to parent genomes and reads counted over SNPs using PAC with the unphased VCF generated above. Read counts were summed across libraries using R and filtered for SNPs with high confidence from WGS (as described above) and high confidence for RNA-seq (> 40 reads in total and > 10 reads coming from either the a or b allele). SNPs were identified as subject to silencing with a log2(abRatio) greater than 1 or less than -1. SNP haplotype was reassigned according to the direction of the log2(abRatio) with positive log2(abRatio) remaining in place and negative log2(abRatio) switching alleles.

### Comparison between cell lines

We used bedtools to intersect a bedfile of SNPs found in each cell line with a bedfile of gene coordinates to assign each SNP to its gene. We then used R to intersect the lists of silenced genes in each cell line.

## Code availability

All scripts and code used in this work is available at Protocols.io https://www.protocols.io/private/44A627E3547B11F1B9DE0A58A9FEAC02

## DISCUSSION

We present here a method to easily and inexpensively identify SNPs within any clonal cell population with more than one X chromosome and assign SNPs within X chromosome genes to the active or inactive copy of the X chromosome. We show how to easily use publicly available haplotype data for commonly used XX cell lines to identify genes within these cells that are subject to XCI silencing. The analysis pipeline that accompanies this article clearly demonstrates how to carry out each of these steps, requiring only a minimal understanding of bioinformatics, making this pipeline accessible to non-bioinformatician bench scientists. This approach can be hypothesis generating i.e. the discovery of silenced genes within a cell population that can be explored through further experimentation, for instance our work showing allele specific changes in DNA methylation in CGIs that contained SNPs. Additionally, this approach can be used to investigate the effects of genetic perturbations, drug effects or other experimental approaches on the levels of XCI silencing in an experimental population, for instance knocking out a key player in XCI and observing changes in expression of silenced genes specifically from the Xi.

We further show how this same principle can be applied to any clonal human cell population with more than one X chromosome through the use of GATK and PAC with WGS and RNA-seq. This allows for investigation of XCI in cell lines that do not have publicly available haplotype data or for which the cost of Hi-C and long read sequencing is prohibitive. This opens the possibility of investigating the effects of patient mutations within cell lines generated from patient samples rather than attempting to replicate the mutations within a more common cell type.

We have found that our methods are able to robustly identify genes subject to XCI silencing in multiple clonal cell populations using both in house generated libraries as well as by leveraging publicly available datasets, opening the possibility to use the wealth of available cell line datasets that are available for further analysis. This method can also accurately identify genes that escape XCI silencing following perturbation of an XCI pathway.

This method cannot be used to identify the Xi or silenced genes in non-clonal cell lines or tissue samples as this method relies on the assumption that the same X chromosome is silenced across the whole population which due to the random nature of XCI is not true in non-clonal populations. This limits this methods useability as it cannot be applied to population datasets or patient samples that were not used to generate clonal cell lines.

## Acknowledgements

We acknowledge that much of the computational work reported in this paper was performed on the Shared Computing Cluster which is administered by Boston University’s Research Computing Services. This work was supported by grants from NIGMS to M.B. (5R01GM144352-03, 5R01GM122893-07). We thank members of the blower lab for critical input on this manuscript.

---

**Supplemental Figure 1:**
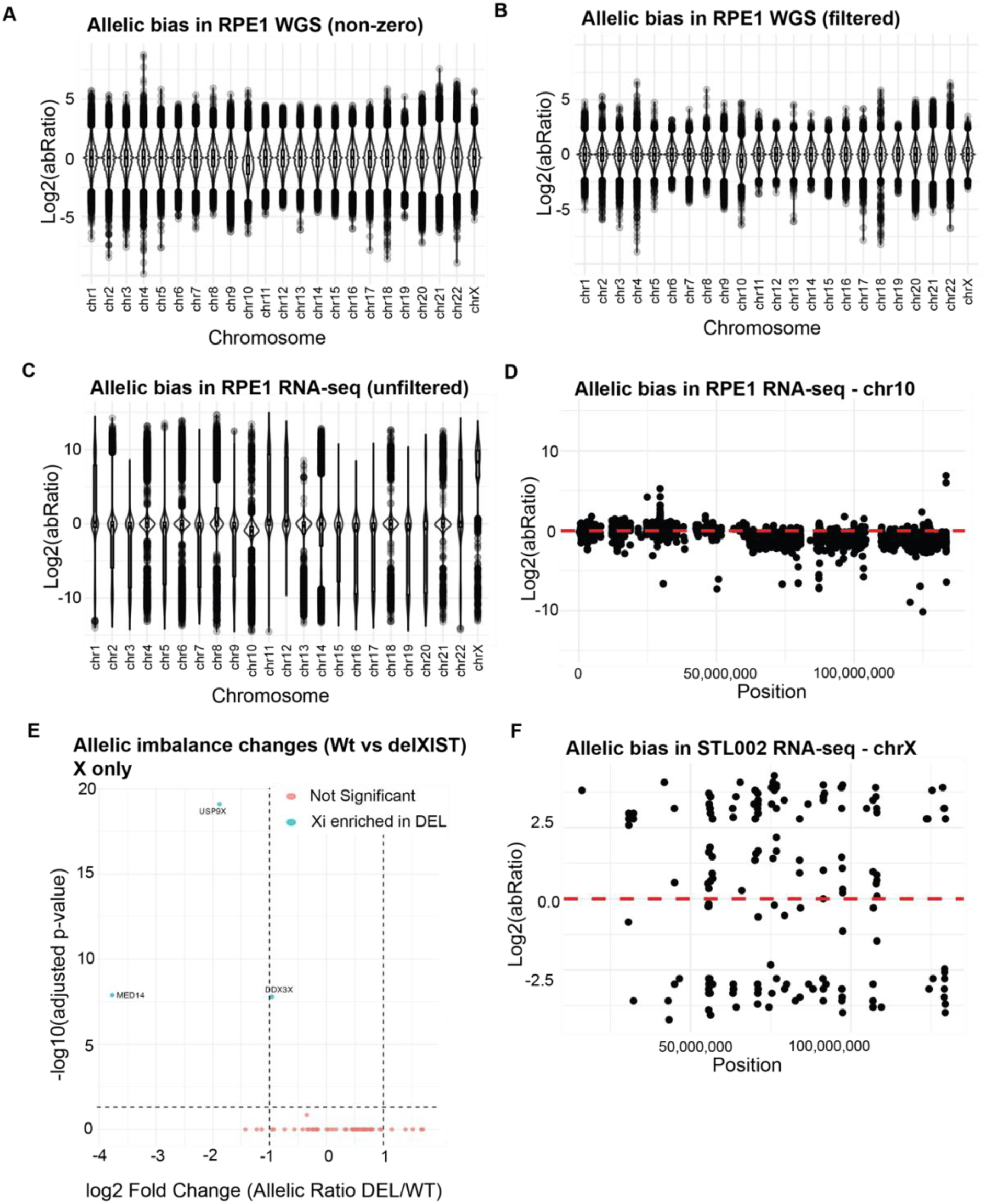
A-B: Allelic bias in WGS of RPE1 for all SNPs with at least 1 read (A) or SNPs with at least 2 reads in allele a, 2 reads in allele b and 5 reads in alleles a+b (B). C: Allelic bias of all SNPs in RNA-seq of RPE1 cells. D: Allelic bias in RPE1 Chromosome 10 showing moderate allelic bias at the site of the Chromosome 10 duplication onto the Xa. E: Volcano plot of log2 Fold Change in the allelic ratio of RPE1 cells with XIST deleted vs WT, showing three genes significantly derepressed by XIST deletion. F: Allelic bias in the X chromosome of RNA-seq from a cell line derived from patient tissue (STL002) showing no clear bias to one X chromosome or the other.

